# From mountaintops to metacollections: using genomics to evaluate ex situ conservation collections. A case study from tropical montane cloud forest plants in Australia

**DOI:** 10.64898/2026.06.27.734930

**Authors:** Manuela Cascini, Lalita Simpson, Stuart Worboys, Warren Worboys, Lydia Guja, Zoe Knapp, Peter Bredell, Julie Percival, Maurizio Rossetto, Darren Crayn

## Abstract

A core aim of ex situ conservation is to represent wild genetic diversity in managed living collections. For the climate-threatened tropical montane cloud forest (TMCF) flora of northeast Australia, an ex situ metacollection of plants and seeds has been established by the Tropical Mountain Plant Science (TroMPS) project. In this study we used reduced-representation sequencing (DArTseq) of wild, herbarium, and ex situ material alongside provenance information for ten species, to pursue two central aims: to characterise landscape-scale genetic structure across species’ ranges, and to evaluate how well the assembled metacollections represent that wild diversity. Analyses revealed consistent patterns of genetic differentiation among mountain top populations across multiple species, reflecting the isolating influence of lowland gaps between upland habitats, with the degree of differentiation varying among species. These results provide the first genetic baseline for Australian TMCF flora and reinforce the importance of treating individual mountain top populations as distinct units for conservation management. Additionally, the project provided valuable insights into the logistical challenges of coordinated multi-institutional collecting, informing strategies for metacollection design more broadly. Evaluation of the metacollection revealed both strengths and gaps in representation across species, providing evidence to refine the current holdings and guide future targeted collecting to strengthen their long-term conservation value.

## Implications for Managers

Genotyping can identify duplicated genets, helping managers optimise space, resources and representation within ex situ collections.

Replicated genets should be intentionally distributed across multiple institutions to reduce vulnerability to loss from local failures.

Ideally, genomic data from wild populations should be generated before field collection to guide sampling and improve metacollection outcomes.

Where pre-collection data are unavailable, retrospective genomic analysis, including herbarium samples where possible, can help evaluate how well ex situ collections capture wild diversity.

## 1 Introduction

With global biodiversity declining at unprecedented rates, ex situ collections represent an important conservation strategy for plants threatened by rapidly changing climates (Pereira et al., 2010; Abeli et al., 2019). These living collections aim to secure species outside their natural habitats, preserving material for future restoration, research, and recovery. A metacollection (sensu BGCI, 2019; Griffith et al. 2020) is an ex situ collection distributed across multiple institutions but managed as a single shared resource, with each institution retaining the plants it grows. Metacollections have emerged as an increasingly valuable approach for improving representation and reducing the risk of loss of ex situ material and maintenance costs, but to achieve these objectives they must be assembled carefully.

Botanic gardens collectively hold over 40% of known threatened plant species, yet their collections are heavily biased toward temperate species, with tropical ones markedly underrepresented (Mounce et al., 2017; Cano et al., 2025). Moreover, species presence in collections does not necessarily equate to adequate representation of genetic diversity, which underpins a species’ evolutionary potential, capacity to adapt or translocation potential (Commander et al., 2018; Hoban 2019). Achieving such representativeness begins with sampling: one must know what and where to sample (Cascini et al., 2025). Ideally, this requires at least some understanding of population genetic structure and access to populations that reflect that structure. Although methodologies for optimising representativeness exist (Bragg et al., 2021; van der Merwe et al., 2021; Fahey et al., 2025; Dimon et al., 2026), their application needs to take into account practical challenges and the immediacy of expected outcomes. Plants may be rare, difficult to access, or seasonally unavailable, and once collected they require ongoing horticultural care and curation. Under such conditions, conservation efforts tend to prioritise maintaining the number of living individuals and may unintentionally do so at the expense of genetic diversity. Consequently, collections assembled opportunistically or founded from few individuals may experience genetic drift, inbreeding, or adaptation to cultivation, ultimately reducing their conservation value (Havens, 2006; Ensslin et al., 2015). Poorly planned metacollections can be low in genetic diversity, overrepresent clonal material, and under-represent adaptive potential. Using such unrepresentative collections for conservation actions (e.g. translocation or population augmentation) may even endanger natural populations via genetic swamping and reduced fitness, rather than enhance diversity through the inclusion of multiple, distinct genets (Bragg et al., 2021; Rossetto et al., 2024). Genetic evaluation can help avoid these issues by identifying redundancy and gaps, improving representation, and providing a strong basis for targeted future collecting and curation with minimal additional investment (Cascini et al., 2025; Doyle et al., 2025; Rossetto et al., 2024).

The Tropical Mountain Plant Science program (TroMPS) is a broad conservation initiative focused on climate-threatened tropical mountain cloud forest (TMCF) flora. The program encompasses several interconnected priorities including ex situ conservation, physiological assessment of climate tolerance, and public engagement. To date, TroMPS research has contributed substantially to the understanding and conservation of Australian TMCF flora, including the description of cryptogam species new to science (e.g. Ramsay et al., 2017; Renner & Worboys, 2018; Meagher et al., 2020), documentation of species previously unrecorded in Australia (Renner & Field, 2019), and the rediscovery of species presumed extinct (Field & Renner, 2019). The program has also generated advances in seed ecology and ex situ management, including new insights into seed dormancy (Hoyle et al., 2023b) and approaches to banking TMCF species (Hoyle et al., 2023a). Central to the program are the establishment of a secure ex situ metacollection and the characterisation of landscape-scale genetic diversity, both to understand how diversity is structured across wild populations, and to evaluate how well the ex situ metacollections represent that diversity to inform ongoing conservation strategies.

In Australia, TMCF is almost entirely restricted to the Wet Tropics of Queensland World Heritage Area (WTWHA). The WTWHA protects 45% of the Wet Tropics bioregion, an area of Australia’s northeast coast defined by very high rainfall, steep north-south mountain ranges and characteristic flora and fauna assemblages. It covers approximately 900,000 ha of predominantly rainforest and is internationally recognised for its outstanding natural value, meeting all four natural World Heritage criteria and representing one of the most complete and diverse living records of the major stages in the evolution of land plants (Wet Tropics Management Authority 2014; Metcalfe & Ford 2008 and Kooyman et al. 2013, UNESCO 1992-2026). It supports more than 3,000 vascular plant species, including over 550 endemic species and 44 endemic genera, which together make up about 18% of Australia’s vascular flora (UNESCO 1992-2026). Topographically, it is characterised by dissected plateaus, steep escarpments, and a patchwork of mountaintops rising above 1000 m. These mountains serve as “sky islands”, refugia of cool, moist forest isolated by “seas” of lowland rainforest. Despite the significant conservation protection of the WTWHA, the high elevation TMCF ecosystems have limited capacity to adapt or shift upslope and consequently are particularly vulnerable to climate change, with projected impacts expected to become evident within this century (Costion et al., 2015; Gempo et al., 2024). This threat is driving ex situ conservation actions, which require an understanding of genetic diversity and structure to enable representative metacollections to be maintained.

Understanding of landscape scale genetic structure within the WTWHA has been shaped largely by studies of vertebrate fauna, which collectively provide a well-resolved framework for interpreting the region’s evolutionary history. Comparative phylogeography of rainforest-dependent reptiles and amphibians has revealed congruent patterns of deep genetic divergence consistent with long-term persistence in at least two major refugial areas separated by the Black Mountain Corridor — a lowland gap in the rainforest that has acted as a persistent phylogeographic barrier (Schneider & Moritz 1999; Schneider et al. 1998). The genetic discontinuities and refugial signatures identified in these studies are consistent with a history of repeated Pleistocene climatic oscillations that drove cycles of rainforest contraction and expansion across the region (Kershaw 1994; Hilbert et al. 2007). Comparable studies of plant taxa have been far fewer, and those conducted on broadly distributed rainforest tree species reveal more variable patterns than those observed for fauna. While some plant species show genetic structuring consistent with co-occurring vertebrate species, including population differentiation associated with key topographic barriers and signatures of refugial persistence, patterns vary considerably among species, apparently reflecting differences in dispersal ecology and life history (Rossetto et al. 2009; Rossetto et al. 2015; Yap et al. 2020). Together, these studies highlight that while topographic complexity and climatic history have shaped genetic diversity across the WTWHA, the degree and nature of that structuring vary considerably among species.

Despite this foundational work, landscape scale genetic structure among Wet Tropics flora, and particularly high-elevation specialists, remains poorly understood. The TMCF flora remains largely uncharacterised, with the few published plant genetic studies having predominantly targeted canopy trees (e.g. *Elaeocarpus* species; Rossetto et al. 2009) and various broadly distributed rainforest species (Yap et al. 2020). These studies have relied primarily on cpDNA haplotype analyses and microsatellite markers, which, while informative, may lack the resolution to detect cryptic lineages or reconstruct fine-scale demographic history. Genome-wide SNP approaches now offer substantially greater power to characterise evolutionarily significant units, intraspecific diversity, and population connectivity. Yet these methods have rarely been applied to endemic Wet Tropics or TMCF flora, leaving a substantial gap in our understanding of the montane plant communities most vulnerable to climate change.

From the 85 Australian TMCF species targeted for ex situ collection within the TroMPS project (Hoyle et al., 2023a Appendix 1), ten species, including five that are Critically Endangered, were selected for genetic analysis (Table 1). Together they represent a taxonomically and ecologically diverse selection of TMCF flora, spanning canopy trees, understorey trees, and shrubs occupying habitats from sheltered montane rainforest to exposed ridgetop boulder fields and stunted heathland. They vary in dispersal ecology, from species bearing fleshy fruits likely moved by frugivorous vertebrates (e.g. *Cryptocarya bellendenkerana*) to species with dry capsules or winged seeds of limited dispersal range (e.g. *Flindersia oppositifolia*), and in pollination biology, with inferred pollinators ranging from generalist insects (e.g. *Eucryphia wilkiei*) to birds (e.g. *Dracophyllum sayeri*). Many exist as small, isolated populations confined to single mountain tops. This diversity of population structure, population size, and species’ biology and ecology can significantly influence genetic diversity and ex situ conservation outcomes, for example by affecting levels of gene flow, population connectivity, and genetic drift (Frankham et al. 2017; Hoban 2019), making this a valuable suite for assessing how geographic isolation, habitat specialisation, and ecological traits interact to shape genetic structure across the Wet Tropics region. The inclusion of these ten species within the TroMPS metacollection program provides a concurrent opportunity to evaluate how well in situ genetic diversity can be captured in ex situ collections assembled in the absence of genetic data and under real-world logistical constraints.

**Table 1.**
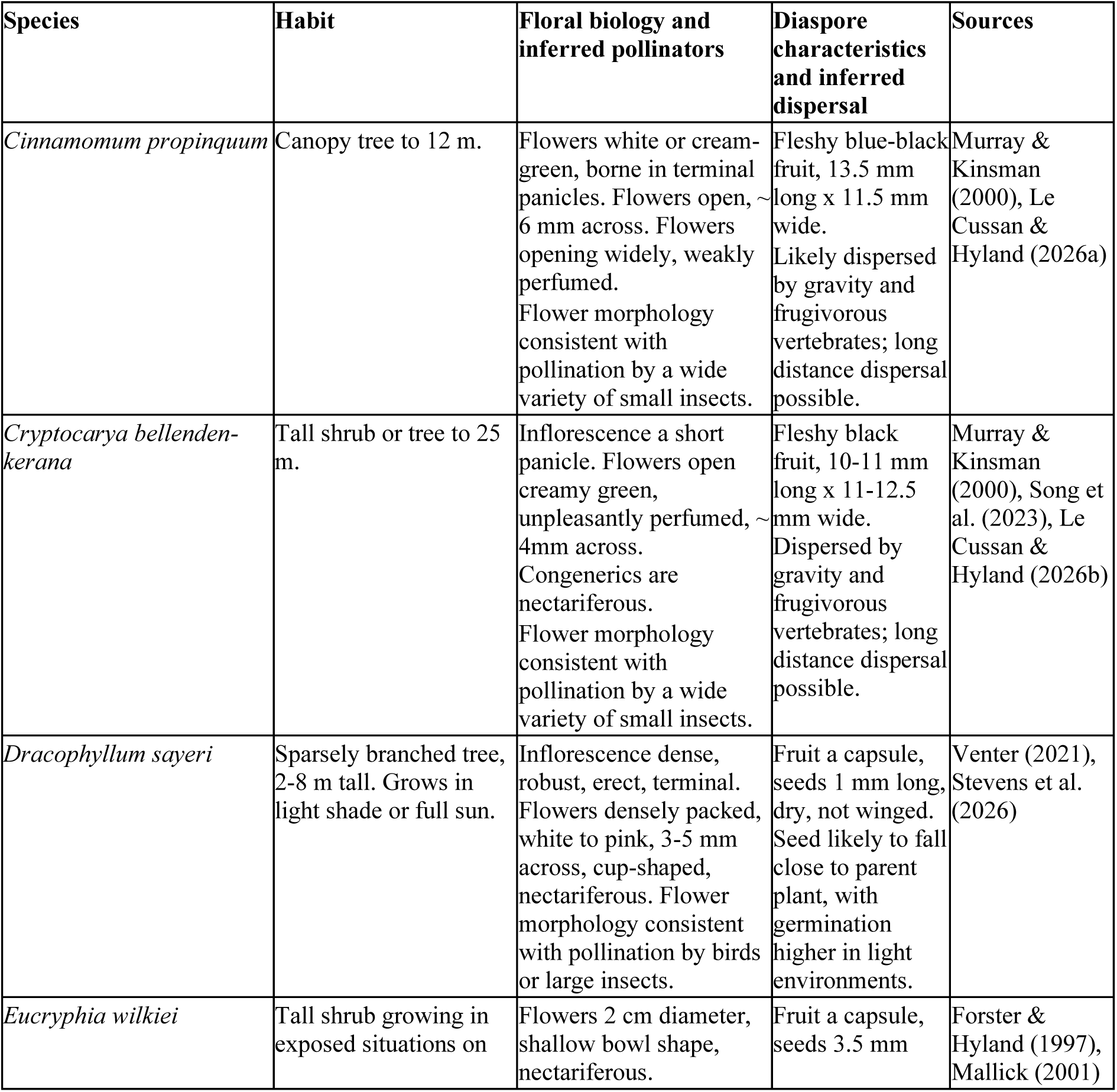
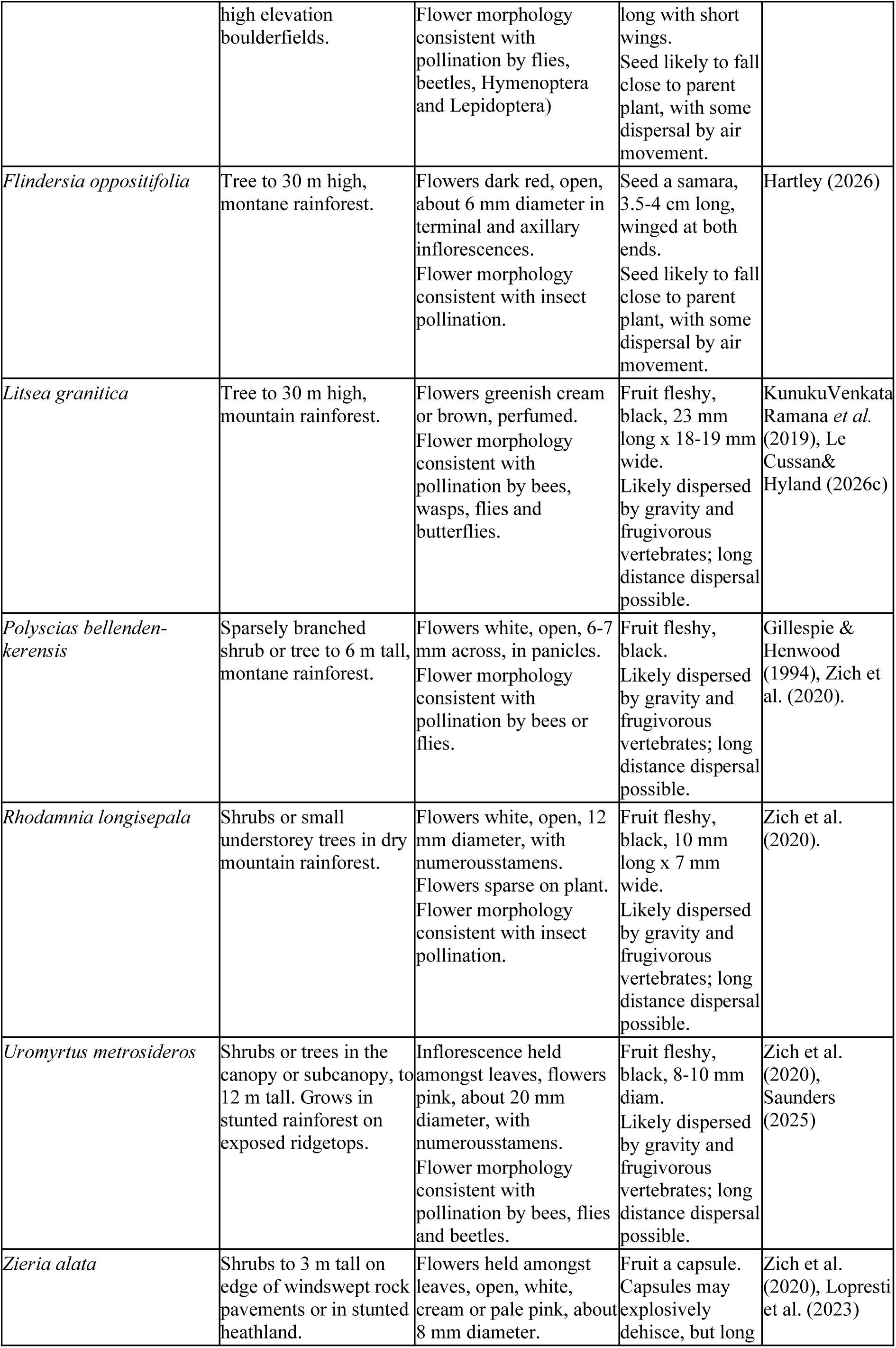
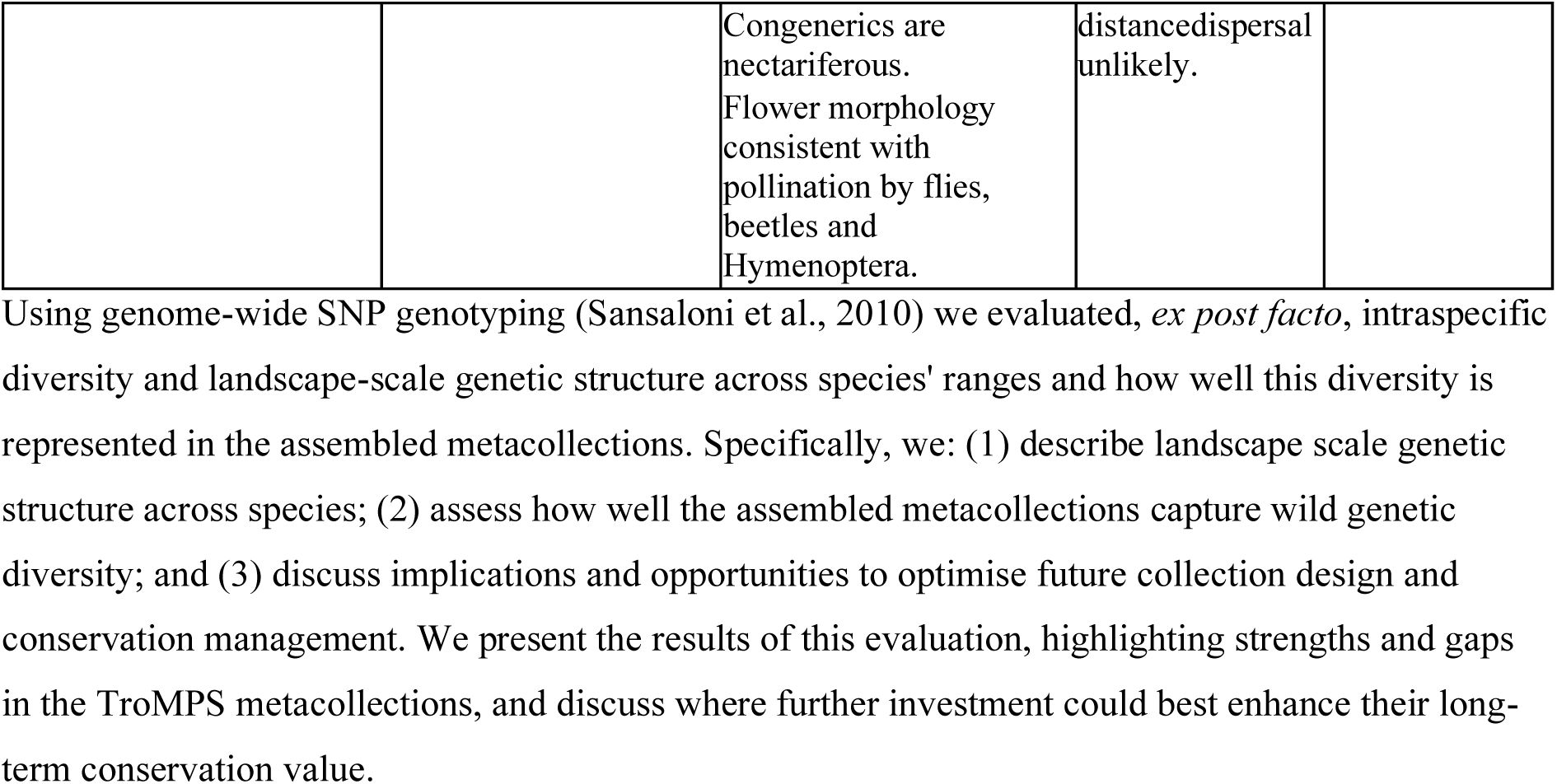
Pollen vectors and seed dispersal ecology inferred from flower and fruit morphology. In the absence of detailed ecological studies, close examination of flower morphology can provide insights into likely pollen vectors (“pollination syndrome” *sensu* Faegri and van der Pijl 1979). Similarly, we can infer likely seed dispersal mechanisms based on fruit morphology. For example, the black fleshy fruit of *Cryptocarya bellendenkerana,* borne in the canopy are likely to attract long-distance dispersers such as top knot pigeons (*Lopholaimus antarcticus*), whereas small white flowers of *Zieria alata* are likely to attract low vagility pollinators such as Diptera and Hymenoptera.

Using genome-wide SNP genotyping (Sansaloni et al., 2010) we evaluated, *ex post facto*, intraspecific diversity and landscape-scale genetic structure across species’ ranges and how well this diversity is represented in the assembled metacollections. Specifically, we: (1) describe landscape scale genetic structure across species; (2) assess how well the assembled metacollections capture wild genetic diversity; and (3) discuss implications and opportunities to optimise future collection design and conservation management. We present the results of this evaluation, highlighting strengths and gaps in the TroMPS metacollections, and discuss where further investment could best enhance their long-term conservation value.

## 2 Methods

For the purposes of this study, an accession was defined as a garden record of material from a field collection, identified by the garden’s accession number and linked to the provenance (mountaintop) of that collection. A single accession may be represented by multiple living plants, and those plants may represent one or more genets. A plant was defined as an individual living specimen maintained in cultivation, and a genet was defined as a genetically distinct lineage identified through genomic analysis. Multiple cultivated plants could therefore represent clonal propagules of the same genet.

### 2.1 Field sampling

Australian tropical montane cloud forest (TMCF) endemic flora were initially identified through an expert workshop held in March 2016, which produced a preliminary checklist of approximately 70 vascular plant species and identified data deficiencies. Targeted field surveys were undertaken in 2016 and 2017 to address these deficiencies, generating more than 850 georeferenced records of target species. These data, together with herbarium records and literature review, were used to finalise a working list of 85 TMCF endemic species (Hoyle et al., 2023a, Appendix 1).

Field collections were conducted between 2016 and 2021 across multiple mountaintops and tablelands within the Wet Tropics World Heritage Area (WTWHA; Figure 1). Collections included pressed herbarium vouchers, silica dried plant tissue for genetic analysis, seed, and vegetative propagation material. Sampling aimed to capture species across their geographic range by collecting from as many occupied mountaintops as possible, with populations defined as plants occurring on a single mountaintop. Representative herbarium vouchers were lodged at the Australian Tropical Herbarium (CNS).

**Figure 1.**
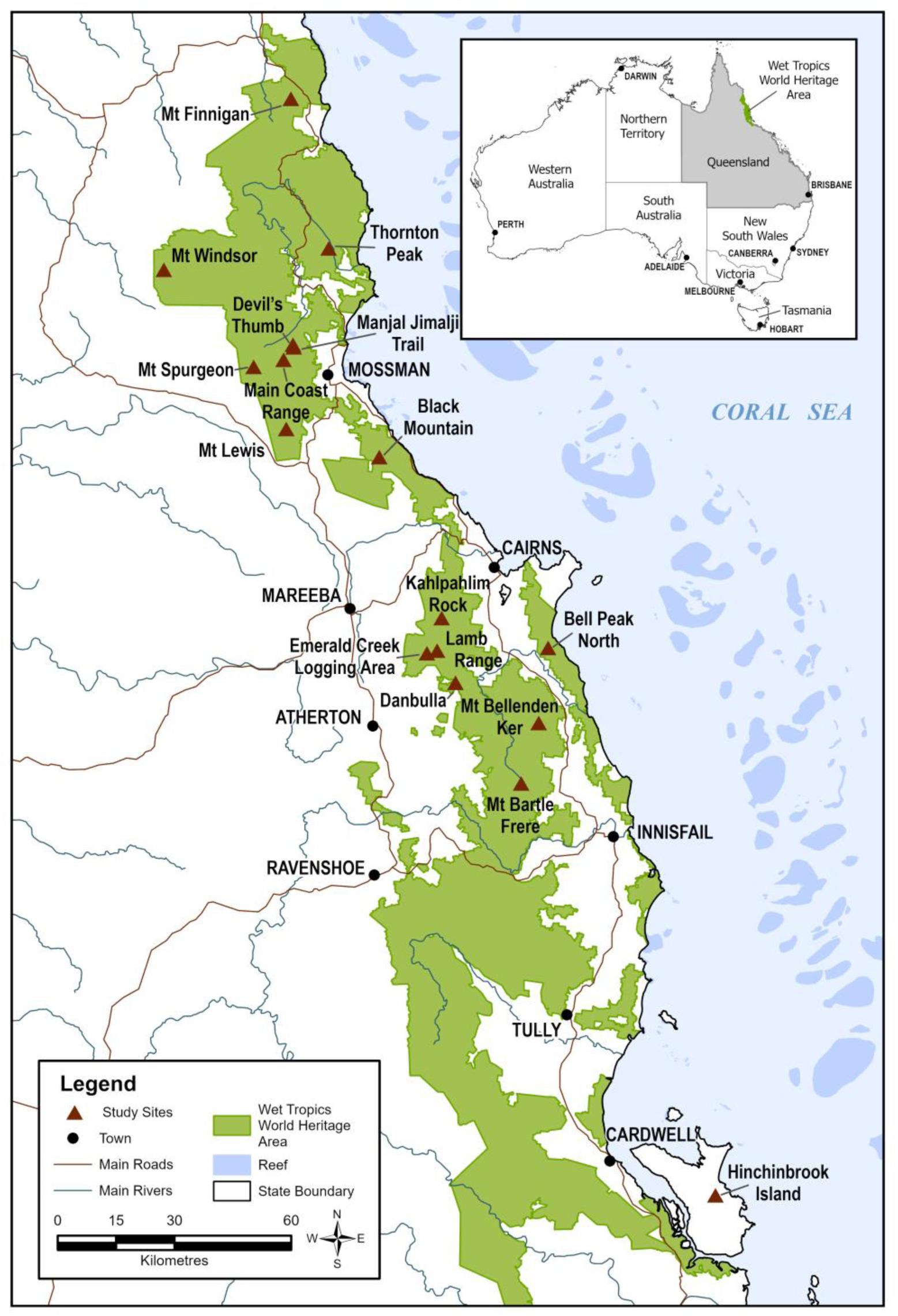
Geographic distribution of principal sampling sites for the ten target tropical montane cloud forest species included in this study. Sites were grouped into populations for analysis: Devil’s Thumb, Mt Spurgeon, Main Coast Range and Manjal Jimalji Trail as Devil’s Thumb–Mt Spurgeon; Kahlpahlim Rock with Lamb Range; Emerald Creek Logging Area with Danbulla; and Bell Peak North as Bell Peak.

For genetic analyses, leaf material was typically collected from five spatially separated plants per population and dried in silica gel. Vegetative material (cuttings and where available, seedlings) were transferred to ex situ conservation facilities at Australian National Botanic Gardens (ANBG) and Royal Botanic Gardens Victoria, Cranbourne (RBGVIC) for propagation and cultivation. As many target species had not previously been maintained in cultivation, propagation methods were refined experimentally, and additional collections were sometimes required. All collected material was accessioned and linked to associated voucher and collection data where possible, allowing provenance to be tracked through propagation, banking, and ex situ cultivation. Of the 583 ex situ plants genotyped, 522 (90%) were linked only to a population voucher (a herbarium specimen from the same mountaintop), 58 (10%; all *Eucryphia wilkiei* and *Rhodamnia longisepala*) to a voucher of their individual source plant, and three to no voucher, so living collections could rarely be traced to individual source plants, reflecting the logistical constraints associated with collecting across remote TMCF populations and maintaining genetically distinct lineages in cultivation. This highlighted the importance of subsequent genetic screening to characterise the diversity captured within ex situ collections.

### 2.2 Plant propagation and distribution

Plant material was propagated using a range of methods tailored to material type and species-specific requirements. Where little prior knowledge existed, approaches were informed by experience with related taxa, and multiple treatments were trialled when sufficient material was available.

A coordinated metacollection was established initially across the ANBG and RBGVIC, and later selected plants were distributed to other participating gardens, e.g. Dandenong Ranges Botanic Gardens. Distributing living collections across multiple institutions provided intentional redundancy to reduce risks associated with maintaining ex situ collections of highly threatened species.

### 2.3 Genetic sampling

Samples for genetic analysis was collected from three sources 1) silica dried tissue collected from established ex situ collections (ex situ); 2) tissue sampled from herbarium specimens (herbarium) and 3) silica dried tissue collected from wild populations during field surveys (wild). Sampling of ex situ propagated material for genetic analyses was undertaken at ANBG and RBGVIC between November 2021 and January 2022, where all propagated plants were held. A unique ID was assigned to each sample at the time of collection, and tissue was dried on silica. Destructive sampling of herbarium specimens was undertaken in the same period. Approximately 20 mg of dried leaf tissue was removed from specimens held at the Australian Tropical Herbarium (CNS) and stored on silica prior to DNA extraction. Silica dried plant tissue collected for genetic analysis during field surveys carried out between 2016 to 2022 was also sampled. All samples (wild, herbarium, and ex situ) are listed in

Supplementary Tables S1. The geographic scope and sampling intensity of samples successfully genotyped for each species is summarised in Table S2.1, and sampling locations for wild, herbarium, and ex situ material are shown in Figure 1. Leaf tissue from wild, herbarium, and ex situ specimens was sent to Diversity Arrays Technology Pty Ltd (DArT, Canberra; Sansaloni et al., 2010) for DNA extraction, library preparation, and DArTseq genotyping following in-house protocols.

### 2.4 Genomic analyses

Loci were retained if they had a reproducibility score of at least 0.96 and no more than 30% missing data; one SNP was kept per sequence tag, monomorphic loci were removed, and samples with more than 50% missing data were excluded (Table S2.2). Samples of other species included in the same DArTseq orders were excluded using the original collection records. Two herbarium specimens whose genotypes identified them as another species were also excluded: one recorded as *Uromyrtus metrosideros* (Kahlpahlim Rock), which was genetically *Uromyrtus tenella* (Section 3.4), and one recorded as *Litsea granitica* (Mt Windsor), which was genetically *L. bindoniana* (Table S2.6).

Samples that could not be distinguished genetically (ramets of the same genet) were identified from two pairwise distances calculated on loci with minor allele frequency ≥ 0.05 and ≤ 20% missing data: the proportion of loci at which the two samples were opposite homozygotes (IBS0), and the proportion of loci with any genotype difference (mismatch). IBS0 is largely insensitive to heterozygote drop-out, the main source of genotyping error in DArTseq data, whereas mismatch separates clones from close relatives such as parent-offspring pairs and full sibs, which share an allele at almost every locus but differ in many genotypes. Kinship estimators assume low genotyping error; in our data, technical replicates of the same sample returned kinship values well below the theoretical threshold for identical samples (0.354; Manichaikul et al. 2010), so kinship was not used. Pairs of samples below species-specific thresholds on both distances were considered ramets of the same genet, and genets were identified as groups of samples connected by such pairs. Thresholds were set from the distribution of pairwise distances for each species and validated with technical replicates (included for eight species; Table S2.3, Figure S2.2). Genet counts were robust to changing both thresholds by ± 25% (Table S2.3), and each of the 12 wild plants that had been sampled two or three times in the field (same collection number) was assigned to a single genet. Genets could not be resolved for *Rhodamnia longisepala* because technical replicates were not distinguishable from other pairs of samples (308 informative loci).

Overall genomic differentiation within species was assessed using two complementary approaches. Both used one representative per genet (all samples for *Rhodamnia longisepala*, whose genets could not be resolved) and loci with no more than 20% missing data and minor allele frequency ≥ 0.05.

First, we performed principal component analysis (PCA) on centred genotypes, with missing genotypes replaced by the locus mean, in R v4.3.3 (R Core Team 2025). Second, we built phylogenetic networks with the NeighborNet algorithm (Bryant & Moulton 2004) from Euclidean distances between genotypes, using SplitsPy (https://github.com/husonlab/SplitsPy), a Python implementation of the SplitsTree methods (Huson & Bryant 2006). Each population was given the same colour and symbol in all figures. To detect misidentified or admixed individuals, each genotyped individual was also compared with the wild genets of every population of its species and with the samples of related species included in the same DArTseq orders; individuals that were genetically closer to another species, intermediate between species, or much more divergent than any population were flagged (Table S2.6). Genetic diversity of wild and ex situ material was compared using one representative per genet (the sample with least missing data), on loci with no more than 20% missing data and minor allele frequency ≥ 0.05; genets detected in both wild and ex situ samples were included in both groups. We calculated observed heterozygosity (Ho), unbiased expected heterozygosity (He; Nei 1978) and allelic richness (Ar) rarefied at each locus to the smaller of the two groups (El Mousadik & Petit 1996), averaged across loci.

### 2.5 Garden records and accession matching

To evaluate whether the TroMPS project established a secure and representative ex situ metacollection, we first collated accession data from the two participating gardens that propagated field-collected material: the ANBG and RBGVIC. This included surveys of records on plant survival and propagation success. Particular attention was given to the ANBG collection because this institution retained original TroMPS accession tags which enabled unambiguous matching between living plants and genetic samples. Provenance for sequenced accessions was assigned using associated population vouchers, which documented the location and identity of source plants. By integrating accession records with genomic data, we determined how many living plants were genotyped, how many genets were represented in the collection, and the provenance of those genets. This, in turn, allowed us to assess mountaintop coverage, identify redundancy, and provide preliminary insights into the security of the metacollection from the available records.

Known wild populations of each species were compiled from the collection localities of herbarium specimens (Australasian Virtual Herbarium, downloaded through the Atlas of Living Australia, https://www.ala.org.au, in 2025) and of the wild samples genotyped in this study, grouped by mountaintop or isolated upland area; records from the Carbine Tableland (Devil’s Thumb, Mt Spurgeon, Main Coast Range and Manjal Jimalji Trail) were treated as one population. Every known population was supported by at least one herbarium specimen; observation records (plot surveys and citizen science) did not add any population and were not used. Each known population was considered as genotyped (at least one wild sample genotyped), sampled ex situ in 2021–22 (at least one genotyped ex situ plant of that provenance) and held in living collections in 2025 (at least one living plant of that provenance at ANBG or RBGVIC). RBGVIC accessions without recorded provenance were counted as living plants but not assigned to a population. Each genet was categorised according to where it was detected: only in wild samples (silica or herbarium), only in ex situ plants, or in both.

To estimate how much wild genetic variation is held ex situ, we used one representative per genet and calculated the percentage of alleles present in the genotyped wild genets that also occur in at least one ex situ genet. We report this for all alleles and for common alleles (frequency ≥ 0.05 in the wild genets); common alleles are reported only for species with at least ten wild genets, because with fewer genets a single copy already meets or exceeds this frequency. To separate the effect of collection size from that of biased collecting, we compared the observed value with that expected for the same number of wild genets drawn at random (mean of 200 draws), where ex situ genets were fewer than wild genets. For species sampled from several populations, we counted the alleles found only in genotyped populations without ex situ genets and the percentage of these held ex situ, and calculated capture for each population, including its private alleles (Table S2.4). Two genotyped individuals that were highly divergent from all other samples of their species, and are probably misidentified, were excluded from capture estimates (*Cryptocarya bellendenkerana* 408393/408418; *Polyscias bellendenkerensis* 409043); their vouchers should be re-examined.

## 3 Results

### 3.1 Dataset composition

Reduced-representation DArTseq generated variable data yield across the ten TroMPS species (see Table S2.2 for sequencing success and quality control (QC) metrics). After filtering, between 22 (*Dracophyllum sayeri*) and 174 (*Cryptocarya bellendenkerana*) samples per species were retained, with post-QC datasets ranging from 1,850 (Zieria alata) to 55,702 (*C.* bellendenkerana) SNPs.

Across species, most genotyped material originated from ex situ collections, with smaller contributions from wild collections preserved on silica or as herbarium specimens (Figure S2.1). Herbarium specimens contributed the fewest samples, and 25 of the 89 genotyped (28%) failed QC, compared with 9 of 734 silica-dried ex situ and wild samples, consistent with degraded DNA in older specimens. Several species, including *Uromyrtus metrosideros* and *Flindersia oppositifolia*, were dominated by ex situ material, whereas others such as *Cinnamomum propinquum* and *Rhodamnia longisepala* incorporated proportionally more wild samples.

### 3.2 Wild populations, genets and alleles held ex situ

Garden surveys in 2025 recorded 253 living TroMPS plants of nine species at ANBG (23 accessions) and 94 at RBGVIC (18 accessions); *Polyscias bellendenkerensis* was absent from both gardens (Table 2). The plants genotyped in 2021–22 came from 24 ANBG and 21 RBGVIC accessions; Table S2.7 lists every accession with its recorded provenance, its provenance according to the genetic data, the number of plants genotyped and the number of genets among them. At ANBG, plants retained their original TroMPS tags and could be matched to genotypes; at RBGVIC they could not, so RBGVIC plants contribute populations (from accession provenance) but not genets. The ten species are known from one to 12 wild populations (Table 2, Figure 2). Living collections in 2025 held plants from all known populations of *Eucryphia wilkiei* and *Rhodamnia longisepala* (each known from a single population) and of *Cinnamomum propinquum*, from two of three populations of *Flindersia oppositifolia*, two of five of *Litsea granitica* and one of two of *Zieria alata*, and from a quarter or fewer of the known populations of the more widespread species (*Cryptocarya bellendenkerana* 3 of 12, *Uromyrtus metrosideros* 3 of 11, *Dracophyllum sayeri* 1 of 4). Some populations sampled ex situ in 2021–22 no longer had living plants in 2025: *C. bellendenkerana* from Mt Bartle Frere, *U. metrosideros* from Mt Bellenden Ker and both sampled populations of *P. bellendenkerensis* (Figure 2).

**Figure 2.**
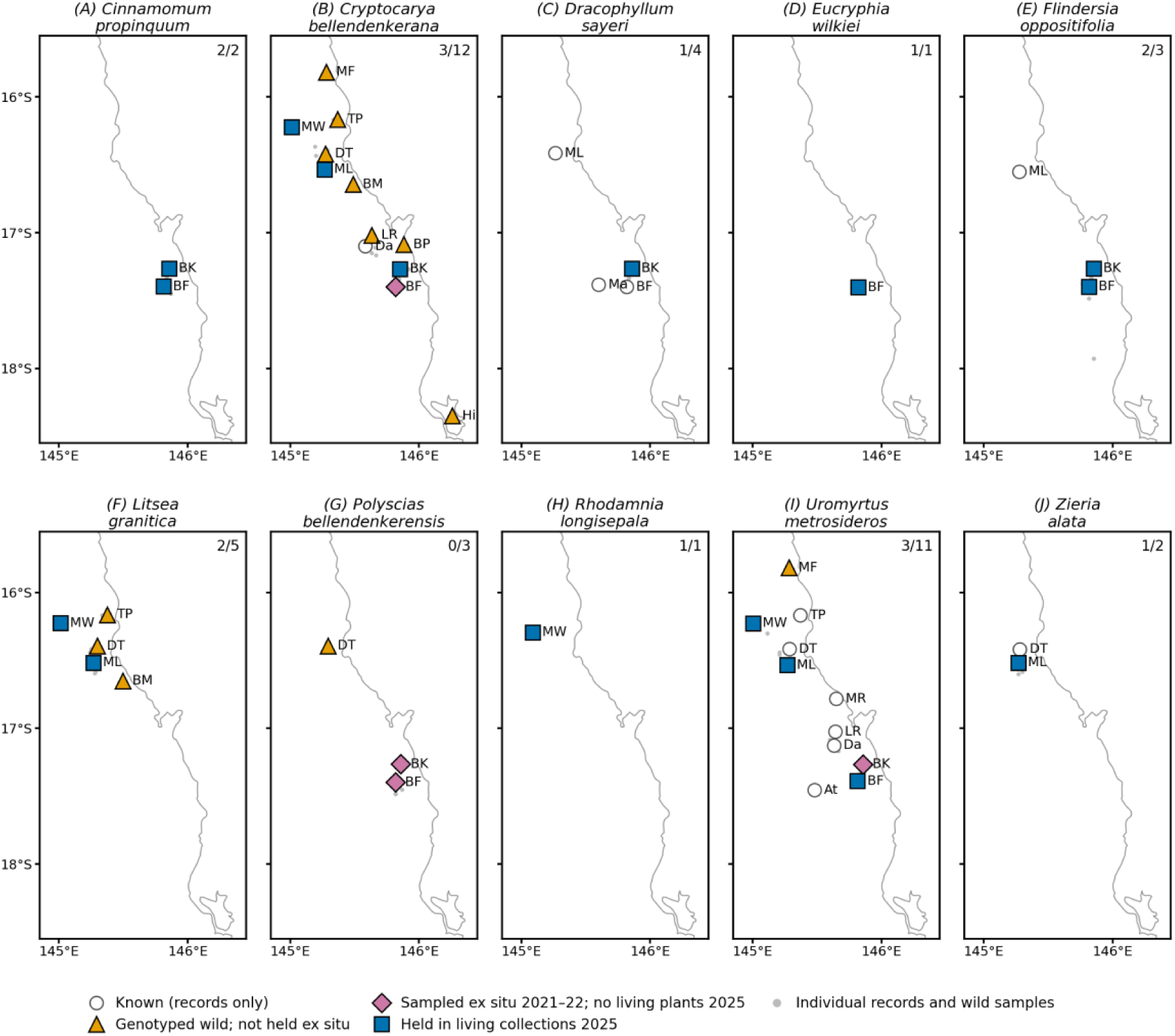
Known wild populations of the ten TroMPS species and how each is represented. Open circles: populations known from herbarium records only; triangles: genotyped wild populations not held ex situ; diamonds: populations sampled ex situ in 2021–22 with no living plants in 2025; squares: populations held in living collections (ANBG or RBGVIC) in 2025. Small grey points: individual herbarium records and wild samples. Numbers in the top right of each panel: populations held in living collections / known populations. BF, Mt Bartle Frere; BK, Mt Bellenden Ker; BM, Black Mountain; BP, Bell Peak; Da, Danbulla; DT, Devil’s Thumb–Mt Spurgeon; Hi, Hinchinbrook Island; LR, Lamb Range; Ma, Malanda; MF, Mt Finnigan; ML, Mt Lewis; MR, Macalister Range; MW, Mt Windsor; TP, Thornton Peak; At, Atherton.

**Table 2.**
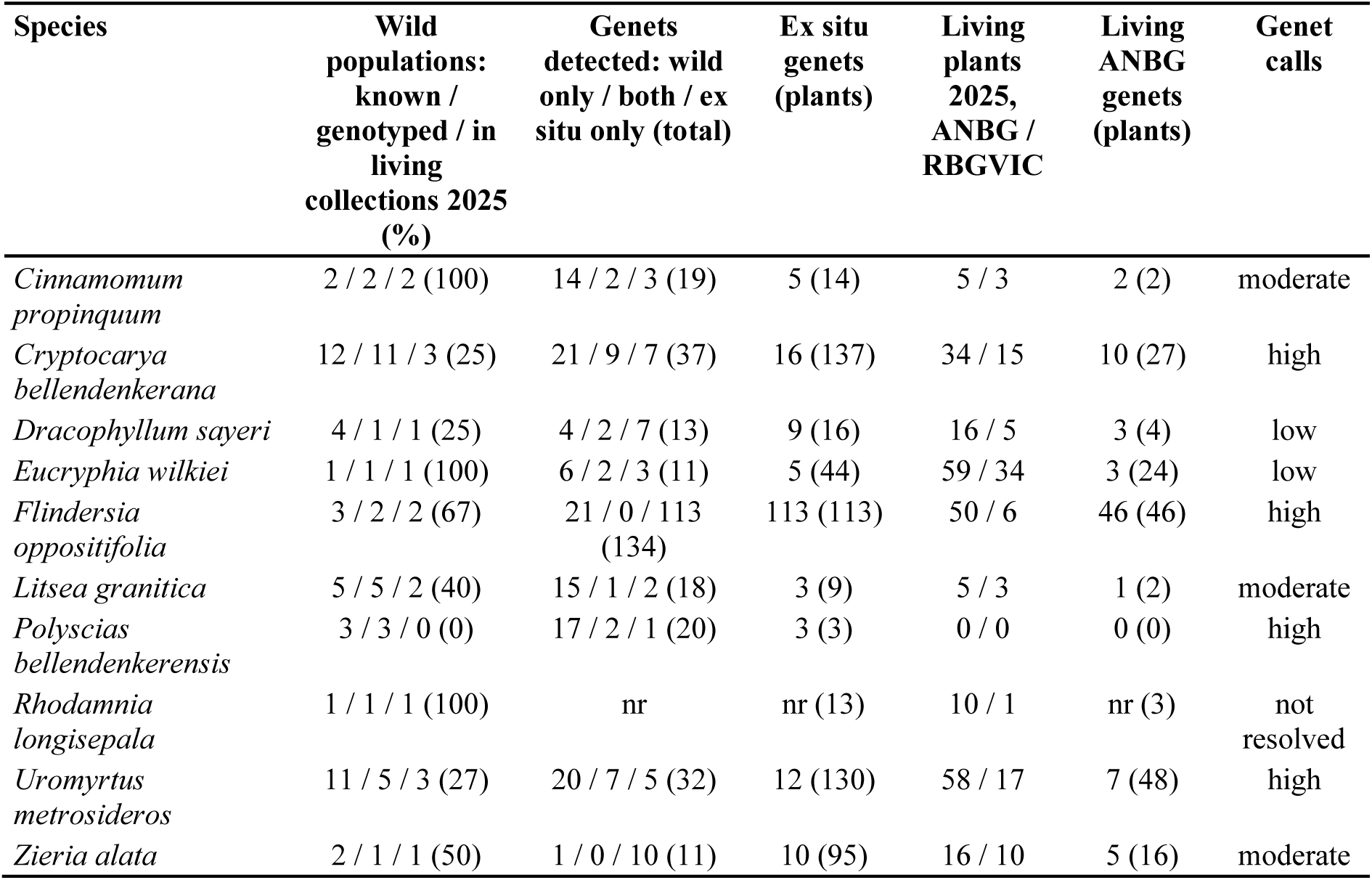
Wild populations and genets of the ten TroMPS species held ex situ. Wild populations: known populations (with at least one herbarium specimen or genotyped wild sample) / populations genotyped / populations represented by at least one living plant at ANBG or RBGVIC in 2025 (percentage of known populations). Genets detected: genets found only in wild samples (silica or herbarium) / in both wild samples and ex situ plants / only in ex situ plants (total; Figure 3). Ex situ genets: distinct genets among the ex situ plants genotyped in 2021–22 (plants genotyped). Living plants: plants alive in the 2025 garden surveys. Living ANBG genets: distinct genets among genotyped plants alive at ANBG in 2025 (plants genotyped); RBGVIC plants could not be matched to genotypes. Genet calls: confidence in genet identification (Table S2.3). nr: genets not resolved.

The number of genets held ex situ was much smaller than the number of plants for most species (Table 2). The 130 genotyped ex situ plants of *Uromyrtus metrosideros* represented 12 genets, the 137 of *Cryptocarya bellendenkerana* 16, the 95 of *Zieria alata* 10 and the 44 of *Eucryphia wilkiei* 5. At ANBG in 2025, the 24 genotyped living plants of *E. wilkiei* represented three genets and the 48 of *U. metrosideros* seven. The exception was *Flindersia oppositifolia*, grown from seed, where each of the 113 plants genotyped was a different genet. Across the nine species with resolved genets, we detected 295 genets: 119 only in wild samples, 151 only ex situ and 25 in both (Figure 3). A genet found in both is a wild individual that was brought into cultivation as cuttings. Genets found only ex situ come mostly from seed-raised collections, in which each seedling is a new genet; all 113 ex situ genets of *F. oppositifolia* are of this kind. Few wild individuals were matched in cultivation (0–9 per species), so whether ex situ collections hold wild variation depends on allele capture rather than on shared genets alone. Clones were also found among wild samples of *E. wilkiei*: on Mt Bartle Frere, three plants collected as separate individuals in 2019 and a herbarium specimen collected about 400 m away in 1986 belonged to one genet, and two further plants collected in 2019 belonged to a second genet. These two wild genets account for 42 of the 44 ex situ plants of *E. wilkiei* genotyped.

**Figure 3.**
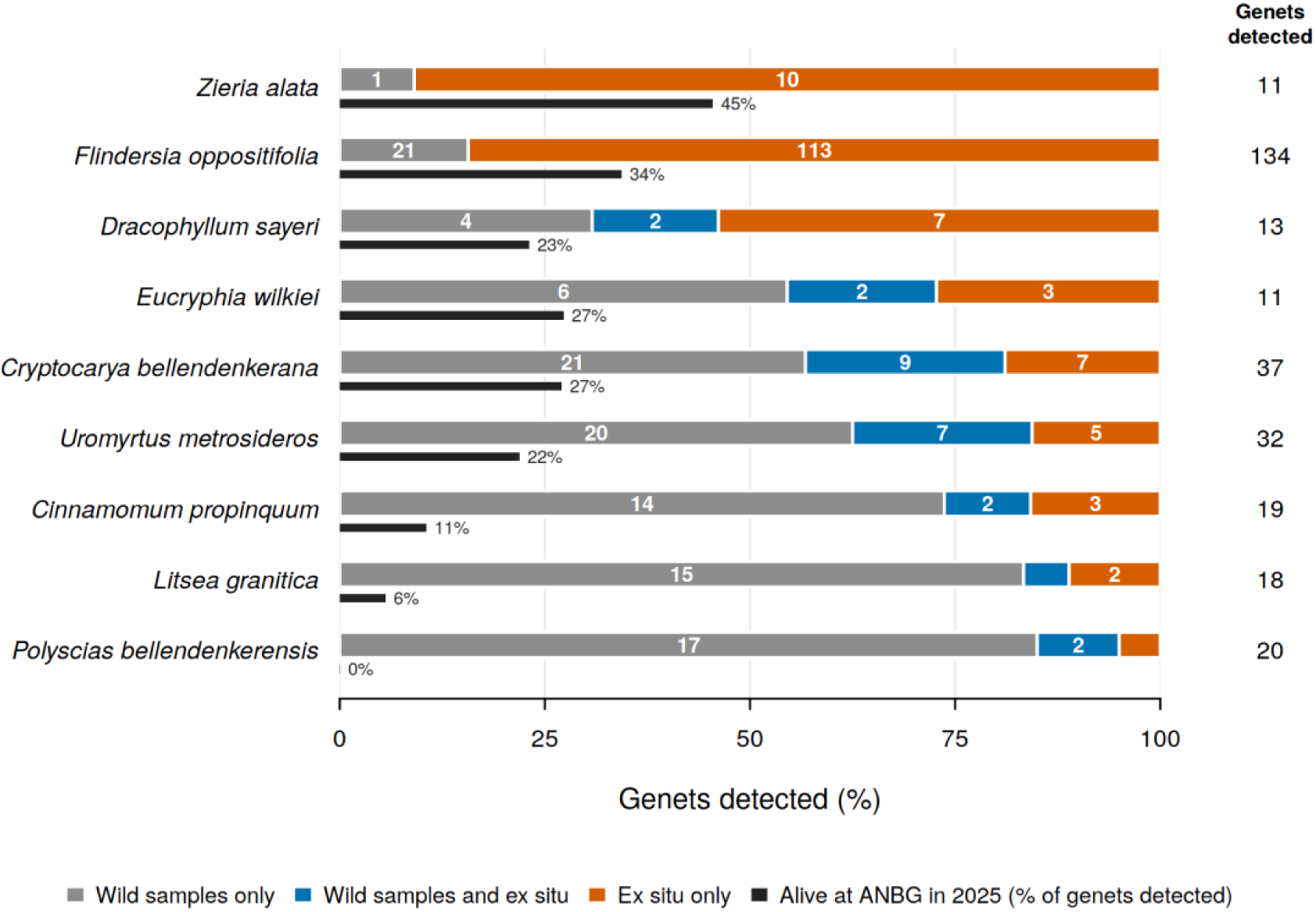
Where the genets detected for each species were found. A genet is a genetically distinct individual; samples whose genotypes matched as clones were treated as one genet (Methods 2.4).Each bar shows how the genets detected for that species were split. Wild samples only (grey): found only in wild samples (silica or herbarium) and matching none of the ex situ plants sampled. Wild samples and ex situ (blue): found in both a wild sample and a garden plant; these garden plants were propagated as cuttings from the sampled wild individual. Ex situ only (orange): found only in garden plants, either grown from seed or propagated from wild plants that were not sampled. Numbers inside the bars are genet counts; the column on the right gives the total genets detected. The thin black bar shows the percentage of genets alive at the Australian National Botanic Gardens (ANBG) in 2025. *Rhodamnia longisepala* is not shown because its genets could not be resolved (Table S2.3). Categories reflect only the plants genotyped: a genet in wild samples only may still be held in ungenotyped garden plants.

RBGVIC accessions without recorded provenance (*Cinnamomum propinquum* 234187, *Litsea granitica* 224529) are counted as living plants but not assigned to any population.

Ex situ genets held most common wild alleles in the species collected from several genets in each of their main populations: 97–99% in *Cryptocarya bellendenkerana*, *Flindersia oppositifolia* and *Uromyrtus metrosideros* (Table 3). Capture was lower where collections derived from few genets or a single population: 83% of common alleles in *Litsea granitica* (three genets, all from Mt Windsor) and 77% of all alleles in *Eucryphia wilkiei*. For *E. wilkiei*, capture was well below that expected from the same number of randomly chosen wild genets (90%), consistent with a collection propagated from a few closely related plants. For *Cinnamomum propinquum* and *C. bellendenkerana*, capture was slightly below the random expectation (72% vs 76% and 91% vs 94% of all alleles); for *L. granitica* and *Polyscias bellendenkerensis* it was within the random range, and for *U. metrosideros* it was above it. For these species, shortfalls mainly reflect the small number of genets collected rather than biased collecting. Capture was lower for the living collections in 2025 than for the plants sampled in 2021– 22, reflecting losses since sampling (e.g. *U. metrosideros* 92% vs 98% of common alleles; *P. bellendenkerensis* 0%, as no plants survive).

**Table 3.** Wild alleles held ex situ. Wild and ex situ genets: genets used for capture estimates (one representative each). Wild alleles held ex situ: percentage of alleles present in the genotyped wild genets that occur in at least one ex situ genet (all alleles / common alleles, frequency ≥ 0.05 in the wild genets; common alleles are shown only for species with at least ten wild genets). Random: expected percentage for the same number of genets drawn at random from the wild genets (mean of 200 draws); not shown where ex situ genets outnumber wild genets. Living ANBG genets: the same for genets alive at ANBG in 2025. Alleles only in populations not collected: alleles present in wild genets from genotyped populations with no ex situ genets and absent from wild genets of the collected populations, with the percentage held ex situ in brackets and the number of such populations in square brackets. *Rhodamnia longisepala*: genets not resolved, so every sample is treated as an individual and values are indicative only. *Zieria alata*: only one wild genet was genotyped, so capture could not be estimated. Two probable misidentifications are excluded (see Methods). Values per population are given in Table S2.4.

| Species | Wild genets | Ex situ genets | Wild alleles held ex situ, all / common (%) | Random, all / common (%) | Living ANBG genets, all / common (%) | Alleles only in populations not collected: n (% held) [populations] |
| --- | --- | --- | --- | --- | --- | --- |
| *Cinnamomum propinquum* | 16 | 5 | 72 / 84 | 76 / 86 | 63 / 75 | – |
| *Cryptocarya bellendenkerana* | 29 | 16 | 91 / 97 | 94 / 99 | 82 / 91 | 4,903 (17) [6] |
| *Dracophyllum sayeri* | 6 | 9 | 93 / – | – | 82 / – | – |
| *Eucryphia wilkiei* | 8 | 5 | 77 / – | 90 / – | 67 / – | – |
| *Flindersia oppositifolia* | 21 | 113 | 94 / 99 | – | 88 / 95 | – |
| *Litsea granitica* | 16 | 3 | 74 / 83 | 74 / 82 | 61 / 69 | 2,491 (17) [4] |
| *Polyscias bellendenkerensis* | 18 | 3 | 79 / 87 | 77 / 85 | 0 / 0 | – |
| *Rhodamnia longisepala* | 24 samples | 19 samples | 56 / 90 | 90 / 98 | 52 / 86 | – |
| *Uromyrtus metrosideros* | 27 | 12 | 94 / 98 | 88 / 98 | 77 / 92 | 493 (3) [1] |
| *Zieria alata* | 1 | 10 | – | – | – | – |

Species-wide capture overstates how well geographically structured variation is held. Alleles found only in genotyped populations with no ex situ genets were largely missing from the collections: 17% of 4,903 such alleles were held for *Cryptocarya bellendenkerana* (six populations not collected), 17% of 2,491 for *Litsea granitica* (four populations) and 3% of 493 for *Uromyrtus metrosideros* (one population, Mt Finnigan; Table 3). By population, 21% of the alleles private to Bell Peak and 8% of those private to Hinchinbrook Island were held for *C. bellendenkerana*, compared with 94% for Mt Bellenden Ker; for *U. metrosideros*, 3% of the alleles private to Mt Finnigan were held (Table S2.4).

### 3.3 Genome-wide variation

Genetic diversity of wild and ex situ genets was similar for most species (Table S2.5). Allelic richness was lower in ex situ than in wild genets for *Rhodamnia longisepala* (1.77 vs 1.97), *Cinnamomum propinquum* (1.62 vs 1.74) and *Litsea granitica* (1.62 vs 1.68), and within 0.04 of the wild value in the other species. Expected heterozygosity was lower ex situ only for *R. longisepala* (0.11 vs 0.17; values based on samples because genets could not be resolved) and differed by less than 0.03 in the other species. *R. longisepala* had the lowest diversity of the ten species. Observed heterozygosity was lower than expected heterozygosity in all species. *Zieria alata* could not be compared because only one wild genet was genotyped.

### 3.4 Landscape-scale genetic structure

Across the ten species studied, PCA and NeighborNet analyses of genome-wide SNP data revealed varied patterns of landscape-scale genetic structure (Figure 4; Supplementary Figures S3.1–S3.9).

**Figure 4.**
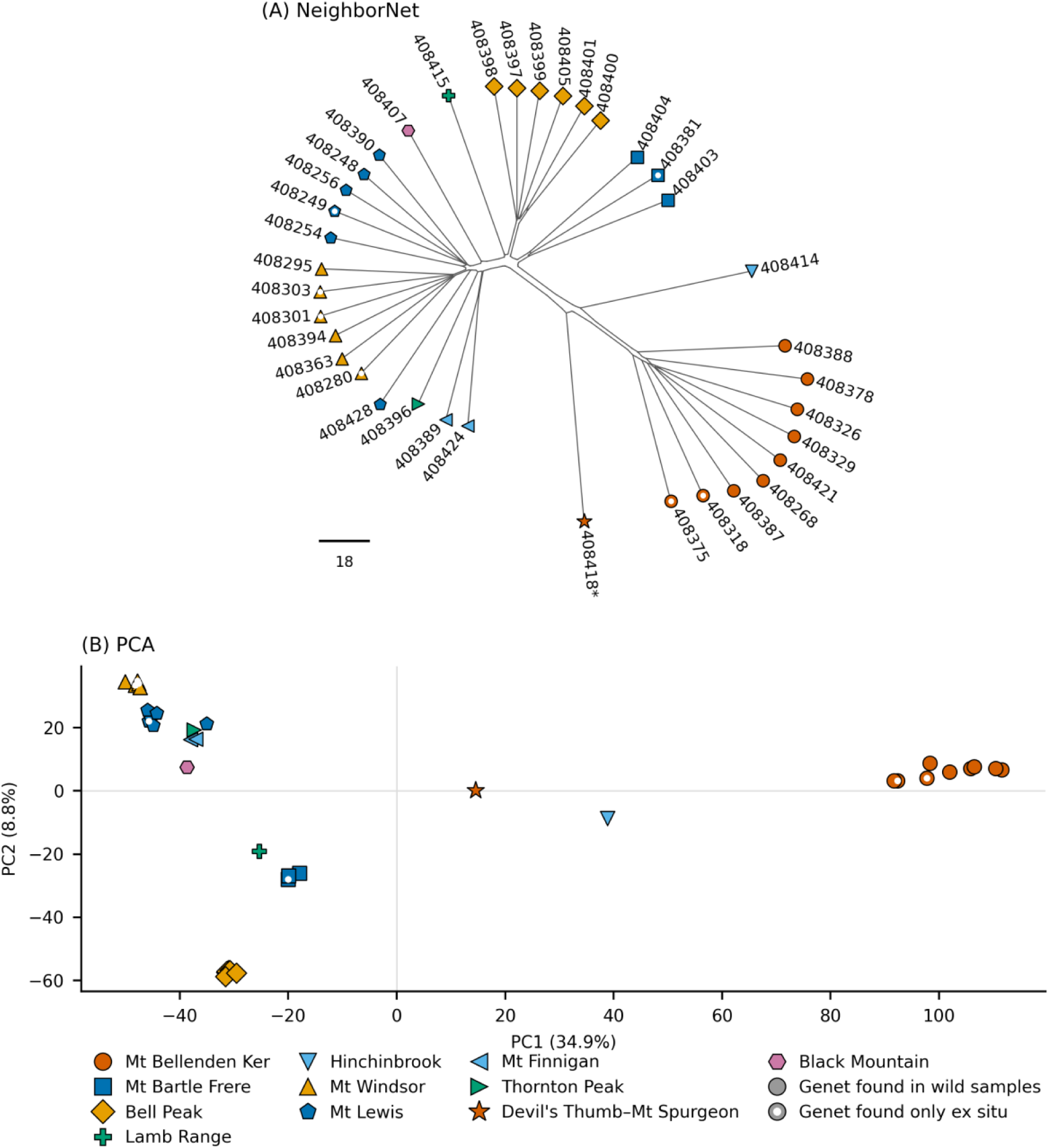
Genomic structure by mountaintop for *Cryptocarya bellendenkerana*. (A) NeighborNet; (B) PCA (PC1 = 34.9%, PC2 = 8.8%); one representative per genet (37 genets, 25,870 loci). Colour and symbol show population and are the same in all figures (Figures S3.1–S3.9); filled symbols are genets found in wild samples and symbols with a white centre genets found only in ex situ collections. *, genet 408393/408418, highly divergent and of uncertain identity (Table S2.6). Scale bar: Euclidean distance. Samples from Mt Bellenden Ker cluster distinctly from other populations, consistent across both analyses.

Among species sampled across multiple populations, analyses showed clear clustering by population. The most consistent cross-species pattern was differentiation between Mt Bellenden Ker and Mt Bartle Frere, evident across all species sampled at both mountaintops, whether the species was restricted to these two mountaintops (*Flindersia oppositifolia*, *Cinnamomum propinquum*) or more widespread across the WTWHA (*Cryptocarya bellendenkerana*, *Uromyrtus metrosideros*, *Polyscias bellendenkerensis*). Despite their geographic proximity as part of the same mountain massif, populations from Bellenden Ker and Mt Bartle Frere formed distinct genetic clusters in all analyses, ranging from strong non-overlap (*F. oppositifolia*) to weaker differentiation (*C. propinquum*).

*Cryptocarya bellendenkerana*, the most broadly sampled species in this study, revealed a hierarchical pattern of genetic structure across the ten populations sampled (Figure 4). Mt Bellenden Ker was the most differentiated population, with a degree of divergence substantially greater than that observed between any other pair of populations. Notably, Mt Bartle Frere clustered within the broader grouping of central WTWHA populations alongside Lamb Range and Bell Peak, showing greater genetic affinity with these geographically distant populations than with the adjacent Mt Bellenden Ker.

Northern WTWHA populations (Mt Windsor, Mt Finnigan, Mt Lewis, Thornton Peak) formed a coherent cluster, and the degree of differentiation between northern and central populations was markedly less than that separating Bellenden Ker from the rest. The Black Mountain individual was intermediate between the northern and central clusters in both analyses, consistent with its transitional geographic position at the Black Mountain Corridor. A single representative from Hinchinbrook Island was also strongly differentiated on a long branch of the NeighborNet and between Mt Bellenden Ker and the other populations on PC1.

Geographic structuring was also evident in the other multi-site species (Supplementary Figures S3.1–S3.9). *Uromyrtus metrosideros*, sampled across five populations spanning both regions, showed clear differentiation between northern and central populations. One Mt Bellenden Ker individual, sampled in the wild and held ex situ as cuttings, was genetically intermediate between *U. metrosideros* and the closely related *Uromyrtus tenella* (Bell Peak), suggesting hybridisation. A herbarium specimen from Kahlpahlim Rock recorded as *U. metrosideros* grouped with *U. tenella* and was excluded from all other analyses of *U. metrosideros*. Within the northern WTWHA, *Litsea granitica* showed clear differentiation between Mt Windsor and Mt Lewis, with the single Thornton Peak and Black Mountain individuals intermediate and the Devil’s Thumb–Mt Spurgeon individual grouping with Mt Lewis. Across the three sampled populations, *Polyscias bellendenkerensis* showed clear differentiation between Mt Bellenden Ker and Mt Bartle Frere, while a single individual from Devil’s Thumb–Mt Spurgeon (Manjal Jimalji Trail) showed higher divergence.

For several species, the results reflect within-population genetic variation only. For *Rhodamnia longisepala* and *Eucryphia wilkiei* this is because both are currently known from a single location, and for *Dracophyllum sayeri* and *Zieria alata* because this study sampled a single site only. Based on current sampling, *R. longisepala* (Mt Windsor) exhibited very low within-population differentiation, while *E. wilkiei* (Mt Bartle Frere) showed comparatively higher variation. *D. sayeri* (Mt Bellenden Ker) and *Z. alata* (Mt Lewis) showed intermediate levels of within-population variation.

## 4 Discussion

In this study, we used genome-wide SNP data (Sansaloni et al., 2010) to retrospectively evaluate the genetic representativeness of the TroMPS metacollectionsof Australian TMCF flora. We combined genomic data, collection metadata from wild samples, and accession records from verified living collections to characterise wild genetic diversity and evaluate how well the assembled collections captured wild variation. This enabled an assessment of how effectively the TroMPS initiative met its goal — to establish a secure, representative ex situ collection — and to identify priorities for further research and targeted conservation action.

### 4.1 Evaluation of the TroMPS metacollections

Genetic diversity measured as allelic richness and heterozygosity was comparable between wild and ex situ populations, indicating that the living collections captured much of the wild variation.

*Rhodamnia longisepala* exhibited the lowest diversity in both wild and ex situ material, consistent with its extremely restricted distribution and Critically Endangered status; the species is known only from a few locations on the Windsor Tableland region of north-eastern Queensland (Snow et al., 2001). Clonality was detected in the wild in *Eucryphia wilkiei*, where plants collected as separate individuals on Mt Bartle Frere, and a herbarium specimen collected there 33 years earlier, belonged to the same genet. This suggests that the species spreads vegetatively in its habitat, so that sampling separate stems in the field does not guarantee distinct genets. It may also explain why the ex situ collection, founded from these two wild genets, holds fewer wild alleles than a random set of the same number of genets (Table 3). Genet calls for *E. wilkiei* are of low confidence (Table S2.3), and the extent of clonality should be confirmed with higher-quality data.

Collection representativeness was mixed, with uneven representation at ANBG and RBGVIC. *Eucryphia wilkiei* is represented by many ex situ plants but at ANBG these are mostly redundant (24 genotyped living plants at ANBG representing three genets). *Uromyrtus metrosideros*, *Flindersia oppositifolia*, and *Cryptocarya bellendenkerana* are comparatively well represented ex situ in terms of both number of individuals and genetic diversity and their ex situ genets held 97–99% of common wild alleles, but they would benefit from broader geographic representation: only 3–17% of the alleles confined to populations not collected were held ex situ. *Dracophyllum sayeri* and *Litsea granitica* are broadly under-represented ex situ. Finally, *Rhodamnia longisepala* is held by few plants (10 at ANBG and one at RBGVIC), whereas *Polyscias bellendenkerensis* is absent from both collections. Many of the study species are novel to horticulture and propagation complexities and constraints limited ex situ establishment success for some species (e.g., *L. granitica*, *Cinnamomum propinquum*, *P. bellendenkerensis* and *R. longisepala* - Scott Levy and Kathryn Scobie pers. comm.).

Mismatches in provenance metadata were also detected: some ex situ plants recorded as originating from one peak clustered genetically with plants from another peak (e.g. 64 *Cryptocarya bellendenkerana*, 14 *Flindersia oppositifolia*, eight *Uromyrtus metrosideros* and two *Cinnamomum propinquum*), demonstrating the importance of accurate recordkeeping, and the value of genomic checks in verifying records. Ultimately, success is measured not by the number of stems grown but by the successful propagation of traceable, representative material that can be effectively used in potential conservation action.

### 4.2 Landscape-scale genetic structure and phylogeographic patterns

The phylogeographic patterns documented here offer the first genetic characterisation of population structure for a suite of Australian tropical mountain cloud forest plant species, a group comprising some of the world’s most range-restricted, evolutionarily significant and climate-threatened flora (Costion et al. 2015; Gempo et al. 2024; Kooyman et al. 2013; Metcalfe & Ford 2008). Despite this significance, these species have been almost entirely absent from the region’s phylogeographic literature, which has focused predominantly on vertebrates with limited evidence from plant taxa (Schneider & Moritz 1999; Schneider et al. 1998; Rossetto et al. 2009; Yap et al. 2020). The most consistent finding, genetic differentiation between populations on Mt Bellenden Ker and Mt Bartle Frere across multiple species, aligns with the model of isolation among upland habitats proposed for the region, in which even short stretches of lowland habitat ecologically unsuitable for upland specialists can function as effective barriers to dispersal and gene flow (Schneider & Moritz 1999; Schneider et al. 1998; Rossetto et al. 2009). Mounts Bellenden Ker and Bartle Frere are among the most climatically stable upland areas in the WTWHA across glacial and interglacial cycles (Hilbert et al. 2007), and despite their geographic proximity, support genetically distinct populations in every multi-site species where both were sampled. These patterns may in part reflect historical refugial dynamics, in which local persistence through repeated Pleistocene climatic contractions established long-term isolation among mountain top populations, with some mountaintops remaining more isolated than others across successive climate cycles (Hilbert et al. 2007; Kershaw 1994; Rossetto et al. 2009; Yap et al. 2020).

The differentiation between northern populations (Mt Windsor, Mt Finnigan, Mt Lewis) and central populations (Mt Bellenden Ker, Mt Bartle Frere) evident in the more widely distributed species *Cryptocarya bellendenkerana*, *Uromyrtus metrosideros* and *Polyscias bellendenkerensis* is broadly consistent with the historical biogeographic framework established for co-occurring fauna, in which the Black Mountain Corridor and associated lowland gaps have been identified as important drivers of genetic discontinuity (Schneider & Moritz 1999). However, the data do not support a simple north- versus-central split cleanly demarcated by the Black Mountain Corridor in the way documented for some vertebrate groups. Rather, genetic differentiation appears to reflect a more continuous geographic gradient, with multiple levels of structure suggesting that the entire series of topographic barriers and lowland gaps throughout the WTWHA contributes to population isolation in plants. In *Litsea granitica*, where Black Mountain was explicitly sampled alongside other northern populations, Black Mountain individuals were somewhat intermediate rather than forming a clearly distinct cluster, consistent with the corridor acting as a transitional zone rather than a sharp genetic boundary.

Variable responses to shared biogeographic barriers have similarly been observed in plant studies from the same region, where species-specific ecology and dispersal history shape the degree of genetic structuring at any given barrier (Rossetto et al. 2009).

While the consistent pattern of divergence among mountain top populations points to geographic isolation as the primary driver of genetic structure in these species, the degree of differentiation varied among species and may in part reflect differences in dispersal ecology. Species bearing fleshy fruits are likely dispersed by frugivorous vertebrates capable of moving seeds considerable distances, whereas species with dry capsules or winged seeds are expected to disperse primarily by gravity or local air movement (Table 1). This distinction is consistent with the observed patterns: *Flindersia oppositifolia*, which produces winged samaras with limited dispersal range, showed the strongest differentiation between Mt Bellenden Ker and Mt Bartle Frere, while *Cinnamomum propinquum*, bearing fleshy fruit, showed weaker differentiation between the same mountaintops. Although these inferences are based on fruit and seed morphology rather than direct dispersal studies, they offer a plausible ecological basis for the variation in degree of genetic isolation observed among species.

A further layer of complexity was evident in *Uromyrtus metrosideros,* where one Mt Bellenden Ker individual was genetically intermediate between *U. metrosideros* and the closely related *Uromyrtus tenella* sampled from Bell Peak, and a herbarium specimen from Kahlpahlim Rock recorded as *U. metrosideros* was genetically *U. tenella*. Whether this reflects ongoing hybridisation, incomplete lineage sorting, or evolutionary lineages that do not align with current species concepts cannot be determined with the current dataset but highlights the importance of comprehensive taxonomic sampling when interpreting genetic structure in species-rich, fragmented floras such as TMCF. A single *Cryptocarya bellendenkerana* individual from Hinchinbrook Island was strongly differentiated from all other populations, suggestive of a genetically distinct peripheral population, though this interpretation warrants caution pending broader sampling.

Within the limits of available sampling and small sample sizes, these results provide both preliminary insight into how diversity is distributed across a poorly known and vulnerable landscape — TMCF — and an important genetic baseline for species that were previously entirely uncharacterised.

### 4.3 Conservation implications and priorities for further action

The congruence in patterns of genetic differentiation documented across multiple species supports treating individual mountain top populations as distinct units for conservation purposes, with important implications both for management of wild populations and ex situ collection design.

Provenance mixing in living collections should be avoided, and any future translocation or augmentation programs should respect population boundaries, as recommended in current guidelines (Commander et al., 2018). Mixing provenances, for example through composite or climate-adjusted provenancing, has been advocated where populations are small or environmental change may outpace local adaptation (Dimon et al., 2026); the differentiation documented here provides the information needed to make that choice deliberately rather than by default. The genetic assessment conducted here also identified specific priorities for strengthening the TroMPS metacollections: (i) propagation trials for difficult species where possible; (ii) targeted wild collecting to fill identified geographic gaps and achieve greater representativeness; (iii) improved accession management through retention of tags/labels, harmonised metadata across institutions, sharing of provenance updates informed by genomic checks, and sharing of relevant protocols; (iv) genotyping of living accessions that currently lack genomic data to better assess completeness. In highly redundant collections, management actions could include either targeted de-duplication to optimise resource allocation or intentional distribution of replicated genets across multiple institutions to reduce vulnerability to catastrophic loss and support evaluation of cultivation outcomes under different horticultural conditions. The findings presented here underscore the value of genetic screening as a tool for guiding metacollection design. Ideally, genomic characterisation of wild populations should be conducted prior to field collecting to better guide and optimise metacollection outcomes (van der Merwe et al. 2023; Cascini et al. 2025; Doyle et al. 2025). However, achieving more comprehensive representation of wild genetic diversity through additional field sampling may be challenging for species distributed across remote, topographically complex, and seasonally inaccessible landscapes, particularly within the logistical and financial constraints typical of conservation programs. In such cases, retrospective genomic assessment, including the potential use of historical herbarium collections where field sampling is not feasible, may provide an important tool for evaluating and improving ex situ conservation outcomes.

Future studies that expand the geographical sampling and statistically assess population structure will be essential to verify the landscape scale genetic patterns identified here and to accurately define the genetically distinct units to guide collection and conservation management priorities. Targeted sampling of under-represented and peripheral populations warrants particular attention, both to resolve ambiguous phylogeographic signals and to fill gaps in ex situ coverage identified by this study. The genetic screen identified two herbarium specimens filed under the wrong species and a probable hybrid between *Uromyrtus metrosideros* and *Uromyrtus tenella* that had been propagated ex situ from cuttings (Table S2.6). Such cases also highlight the importance of verified taxonomic identification in both field collecting and accession management and broader taxonomic sampling of closely related species.

For conservation outcomes, broader sampling of wild populations and better management of ex situ accessions are likely to return more than deeper sequencing. Nevertheless, while the present study is based on reduced-representation genomic data, future expansion to whole-genome sequencing (WGS), which captures variation across the entire genome (Ellegren 2014), would enable substantially deeper inference of evolutionary and demographic processes. Genome-wide data would facilitate more robust reconstruction of historical population size changes, divergence timing, and connectivity among refugial lineages. In addition, WGS would allow more detailed assessment of inbreeding patterns through the analysis of runs of homozygosity and the distribution of potentially deleterious variation (Chen et al. 2025). Such information would provide a stronger evolutionary baseline for evaluating genetic risk, prioritising lineages for conservation, and informing any future management interventions, including augmentation or assisted gene flow.

To synthesise these findings in a conservation management context, the major strengths, gaps, and inferred priority actions identified for each species are summarised in Table 4.

**Table 4.** Summary of population genetic structure, ex situ representation, and conservation priorities for ten TroMPS species. Wild populations: known / genotyped / held in living collections in 2025 (percentage of known populations), as in Table 2. Population structure is based on SNP genotyping; ex situ representation reflects the proportion of known wild populations represented in living metacollections at the time of assessment. Redundancy refers to the ratio of living plants to inferred genets. Priority actions are ranked relative to the severity of representation gaps identified.

| Species | Wild populations: known / genotyped / in living collections 2025 (%) | Population structure | Ex situ representation | Redundancy / representation issue | Main conservation gap | Priority action |
| --- | --- | --- | --- | --- | --- | --- |
| *Cinnamomum propinquum* | 2 / 2 / 2 (100) | Weak but consistent differentiation between Mt Bellenden Ker and Mt Bartle Frere | High geographic representation (100% of populations represented ex situ) but few genets maintained | Only 2 genets represented among 5 ANBG plants | Low effective genet representation within living collections | Increase the number of distinct genets maintained ex situ |
| *Cryptocarya bellendenkerana* | 12 / 11 / 3 (25) | Very high geographic | Moderate–low (11 of 12) | Multiple genets | Incomplete capture of | Prioritise recollecti |
|  |  | structure across WTWHA; Bellenden Ker highly differentiated | populations genotyped but only 25% held in living collections) | maintained ex situ (16 genets; 10 living at ANBG), but many differentiated populations absent from collections | geographically structured diversity | on from underrepresented populations and regions |
| *Dracophyllum sayeri* | 4 / 1 / 1 (25) | Uncertain; single sampled population and low SNP recovery | Low (25% of populations represented ex situ) | Only 3 genets among 4 genotyped living ANBG plants (16 plants alive); genomic resolution limited | Restricted geographic representation and limited genomic resolution | Expand sampling and improve genomic characterisation |
| *Eucryphia wilkiei* | 1 / 1 / 1 (100) | Low geographic structure (single known population) | High geographic representation (100%) | Extremely high redundancy (24 genotyped living ANBG plants correspond to only 3 genets) | Very low effective genetic representation despite many living plants | Prioritise recollection of additional genets, genotyping wild plants first because separate stems may be clones |
| *Flindersia oppositifolia* | 3 / 2 / 2 (67) | Strong differentiation between Mt Bellenden Ker and Mt Bartle Frere | High (67% populations represented ex situ) | Very low redundancy (46 genotyped living ANBG plants = 46 genets) | Some geographically differentiated populations remain absent | Maintain existing collections and expand to additional populations |
| *Litsea granitica* | 5 / 5 / 2 (40) | Moderate differentiation among mountain tops | Low (40% of populations held in living collections) | Only 1 genet among living ANBG plants; 3 ex situ genets in total, all from Mt Windsor | Poor geographic representation in living collections | Expand ex situ collections across additional populations |
| *Polyscias bellendenkerensis* | 3 / 3 / 0 (0) | Strong differentiation between mountaintops | Very low (0% represented ex situ) | No living ex situ collections currently maintained | Complete absence from living metacollections | Highest priority for recollection and establish |

|  |  |  |  |  |  | ment of<br>ex situ<br>collection<br>s |
| --- | --- | --- | --- | --- | --- | --- |
| *Rhodamnia longisepala* | 1 / 1 / 1<br>(100) | Low within-population<br>differentiation (single<br>known<br>population) | Single known<br>population held<br>(10 ANBG<br>plants, 1<br>RBGVIC plant) | Redundancy<br>unknown;<br>genets could<br>not be<br>resolved | Low genetic<br>diversity and<br>limited<br>security of<br>living<br>collections | Maintain<br>collection<br>s and<br>increase<br>genet<br>represent<br>ation<br>where<br>possible |
| *Uromyrtus metrosideros* | 11 / 5 / 3<br>(27) | Strong<br>north–<br>central<br>differentiation; evidence<br>of possible<br>hybridisation | Low–moderate<br>(27%<br>populations<br>represented ex<br>situ) | Several<br>distinct<br>genets<br>maintained<br>ex situ (12<br>genets; 7<br>living at<br>ANBG), but<br>many<br>differentiated<br>populations<br>absent | Incomplete<br>capture of<br>geographical<br>ly structured<br>diversity | Expand<br>collection<br>s across<br>genetically<br>differentiated<br>populations |
| *Zieria alata* | 2 / 1 / 1<br>(50) | Uncertain;<br>only one<br>sampled<br>population | Moderate (50%<br>populations<br>represented ex<br>situ) | Moderate<br>redundancy<br>(16<br>genotyped<br>living<br>ANBG<br>plants<br>correspond<br>to 5 genets) | Limited<br>geographic<br>and genet<br>representation | Increase<br>represent<br>ation of<br>distinct<br>genets<br>and<br>populations |

## 5 Conclusion

As a coordinated, multi-species conservation program sampling across the same mountain top landscapes, the TroMPS initiative provided a unique opportunity to characterise landscape-scale genetic structure across a suite of co-distributed TMCF species. This revealed consistent patterns of differentiation among populations that reflect the isolating influence of lowland gaps between upland habitats, with the degree of differentiation varying among species. Together these findings reinforce the value of treating individual mountain top populations as distinct units for conservation management and provide an important genetic baseline for species that were previously entirely uncharacterised, highlighting the breadth of diversity that must be considered when designing ex situ collections for this flora.

With the establishment of ex situ metacollections a central aim of the program, TroMPS provided a valuable opportunity to test and refine the logistics of coordinated collecting, propagation and data management across institutions, and demonstrated that even retrospective genomic assessment can meaningfully strengthen metacollection outcomes. For a flora that is both evolutionarily irreplaceable and acutely vulnerable to climate change, the steps taken now to build representative, well-documented collections will have lasting conservation value.

